# Annual ploughing turns wildflower plantings into ecological traps for ground-nesting wild bees

**DOI:** 10.64898/2026.08.25.746958

**Authors:** Christopher Hellerich, Alexandra-Maria Klein, Michael Garratt, Anne-Christine Mupepele, Felix Fornoff

## Abstract

Wildflower plantings are an important conservation measure for supporting wild bee diversity. They aim to enhance floral resources for nutrition, but do not explicitly consider that bees also require suitable nesting and overwintering resources. Wildflower plantings may provide nesting habitat for ground-nesting bees, but the effects of soil management, e.g. ploughing, on ground-nesting bees are hardly known. To study how ploughing and age of wildflower plantings affect ground-nesting bees, we sampled bees on ploughed and unploughed wildflower plantings aged 0-4 years. We used emergence traps to sample bees directly after emergence from the ground, allowing inference on nest numbers. We found that ploughing and wildflower planting age negatively affected overwintering bee nest numbers. Nesting peaked in the year of wildflower planting establishment and declined thereafter, indicating lower habitat suitability at later successional stages. Annual ploughing caused the greatest reduction in nest numbers (−72 %), providing evidence for an ecological trap.

## Introduction

Wildflower plantings are a popular tool for short-term, effective nature conservation and ecological restoration (Haaland et al. 2011; von Königslöw et al. 2022; Scheper et al. 2013). Under the European Common Agricultural Policy, they are promoted through both annual (‘eco-schemes’) and perennial (‘agri-environment-climate commitments’) schemes (Šajn 2024). If well implemented, wildflower plantings provide a continuous supply of floral resources throughout the season (Schmidt et al. 2022), which is particularly important for wild bees (Hymenoptera: Anthophila) due to their reliance on nectar and pollen for survival and reproduction (Michener 2007). It has been shown that wildflower plantings can effectively promote wild bees and other pollinators (Grass et al. 2016; von Königslöw et al. 2022; Ouvrard et al. 2018; Williams et al. 2015), as well as other functional groups such as natural enemies (Blaauw & Isaacs 2015), thereby directly contributing to the provision of associated ecosystem services (Albrecht et al. 2020).

Like many other flying insects, most wild bees spend the majority of their lives belowground (estimates range from 64 to 83%) and depend on suitable substrate for nesting, development, and overwintering (Antoine & Forrest 2021). Some studies have demonstrated the value of continuous wildflower plantings for arthropod nesting and overwintering (Boetzl et al. 2022; Ganser et al. 2019), but research focusing specifically on wild bees remains scarce, because studying them during their ground phase is challenging: It is still largely unknown where bees actually nest or overwinter, and the fact that they may use different locations for nesting (i.e., the place of reproduction) than for overwintering makes matters even more intricate (Danforth et al. 2019; Yanega 1990). Because wild bees are central-place foragers, it is efficient for them to nest as close as possible to their foraging resources (Zurbuchen et al. 2010). Indeed, what evidence exists suggests that wildflower plantings do provide nesting and overwintering habitat for wild bees (Williams et al. 2024). Moreover, many species are philopatric and tend to nest near to the mother nest (Danforth et al. 2019), which should lead to increasing nest numbers or aggregations as wildflower plantings age.

However, as factors known to affect bee nesting, such as soil properties and ground cover, may change over time (Antoine & Forrest 2021; Cane 1991; Tsiolis et al. 2022), so may the suitability of wildflower plantings for wild bee nesting and overwintering (Boetzl et al. 2022; Williams et al. 2024). Furthermore, ploughing of the soil during nesting and overwintering may destroy bee nests and thus reduce bee survival but may also alter the site’s attractiveness for nesting in the following season (Cuminale et al. 2026; Shuler et al. 2005; Tschanz et al. 2024; Ullmann et al. 2016; Williams et al. 2010). Ploughing, i.e. the turnover of soil by heavy machinery, is a common farm management practice that has been shown to negatively affect ground-nesting flying insects, with a disproportionate effect on larger species (Hellerich et al. 2026). It usually affects the upper soil layer to a depth of around 20-30 cm and therefore the typical nesting depth of most ground-nesting bees (Harmon-Threatt 2020).

Despite wildflower plantings being widely promoted as a key tool for bee conservation, important knowledge gaps regarding their use as nesting and overwintering habitat and the effects of ploughing remain. To address these gaps, we hypothesized that a) ploughing destroys bee nests and thus negatively affects overwintering bee nest numbers, and that b) smaller bees are less affected by ploughing than larger bees, as it was shown for other soil organisms (Hellerich et al. 2026; Kladivko 2001; Wardle 1995). Leaving wildflower plantings unploughed for several years (e.g., five years under German agri-environment-climate commitments) may provide undisturbed nesting and overwinter habitat (Albrecht et al. 2020), and we thus further hypothesize that c) the number of ground-nesting bee nests increases with the age of unploughed wildflower plantings.

Using soil emergence traps in wildflower plantings in a replicated two-year field experiment, we were able to study bee nesting and the effects of ploughing by quantifying overwintering bee nests across newly established and one-to four-year-old wildflower plantings. Our findings reveal that perennial wildflower plantings provide attractive nesting habitat mostly in the year of establishment and lose attractiveness as they age. We further demonstrate the negative effects of ploughing on bee nests and conclude that through ploughing after one flowering season, annual wildflower plantings can act as ecological traps for ground-nesting wild bees.

## Materials and methods

### Experimental design

Our study sites were twelve wildflower plantings on three conventional farms in the Upper Rhine Valley, southwest Germany (47° 59’ N, 7° 43’ E, 210 mASL). Wildflower plantings varied in age, with seven wildflower plantings established 1-3 years prior to the start of the experiment and five wildflower plantings newly established in spring 2023 at the start of the study. Sampling these plantings over two consecutive years (2023-2024) therefore covered planting ages from 0 to 4 years since establishment, across which we measured the effects of ploughing and planting age on overwintering bee nests. Wildflower plantings contained unploughed control plots and adjacent ploughed treatment plots, dividing the full length of the planting laterally, allowing direct comparisons of the effects of ploughing within the same wildflower planting. An additional control plot was established in 2024 on four wildflower plantings by laterally dividing the existing control plot and data from both plots were included in the analysis. On one wildflower planting, no ploughing treatment could be conducted, and plots thus only served as control plots. Establishment of wildflower plantings and spring ploughing of treatment plots followed local farm practices (mouldboard ploughing to 20-25 cm depth and sowing with a power-harrow seed drill) and control plots remained unploughed throughout the study. This experimental design created a full factorial design combining wildflower planting age and soil disturbance via ploughing, allowing to assess the use of wildflower plantings as nesting and overwintering habitat by ground-nesting wild bees, and the effects of ploughing on bee emergence, across different wildflower planting ages (further details are provided in Hellerich et al. 2026)

### Wild bee sampling

Custom-designed emergence traps (‘e-traps’) were used to sample wild bees directly after ground emergence (see Hellerich et al. (2025) for a detailed description of the e-trap design). This approach allowed collected individuals to be assigned to their nesting place, as the traps prevent capture of bees originating outside the area of ground which they cover. All sampling was conducted with permission from the relevant local authorities.

Eight e-traps were deployed at regular intervals along two transects per plot, each spanning the full length of the wildflower planting (Hellerich et al. 2026). The first trap of each transect was positioned 3 m from the plot edge. Traps remained at the same position throughout each sampling year and were shifted 2 m laterally from their previous year’s location in 2024. Each trap covered 2.2 m^2^, resulting in a total sample area of 17.6 m^2^ per plot. A total of 368 e-traps were deployed on 0-4 year old wildflower plantings (160 traps in April-May 2023 and 208 traps in March 2024). Traps remained active for an average of 99 days (standard deviation = 19) in 2023 and 128 days (standard deviation = 1) in 2024. They were emptied every two weeks, which represented one sampling round. All collected bees were dry-mounted and identified to species level using standard classification keys (Amiet et al. 2017; Schmid-Egger & Scheuchl 1997).

### Bee classification, nest number estimation and body size measurement

For reasons of simplicity, we use the term ‘nest’ in our study as an umbrella term for places of reproduction and of overwintering - the latter being not always the place of reproduction (Yanega 1990). To assess the effect of ploughing on the nests of overwintering bees, only individuals that had emerged from overwintering were included in the analysis, excluding actively nesting bees (i.e., from nests built previously in the same season). To identify whether a bee was overwintering, bees were classified by assessing mandible and wing wear, life history traits such as the level of sociality and the life stage during overwintering, and emergence phenology, to determine the nesting activity of each individual and the number of nests per e-trap sample (Hellerich et al. 2025). Mandible and wing wear were visually assessed using a stereo microscope (Leica S9E). Species’ life history traits and phenology were determined from the literature (Westrich 2018). All bees that could not be clearly attributed to a nesting activity were classified as ‘unknown’ and excluded from analysis.

The number of nests was used as response variable in the statistical analysis (see Hellerich et al. 2025). This approach was chosen to minimize biases associated with differing life history traits among species. Using the number of bee individuals as response variable could distort results, first because the level of sociality influences the number of offspring per nest (Cuminale et al. 2026), and second because worker generations of eusocial species do not reproduce. Although such biases are reduced when studying overwintering bees, for example because commonly worker generations do not overwinter, using nest counts still offers a more standardized and comparable approach (Hellerich et al. 2025).

For testing whether ploughing disproportionally affects larger bees, intertegular distance (‘ITD’) of all overwintering bees was used as an indicator of body size (Cane 1987). The inbuilt ocular micrometer of the stereo microscope was used to measure ITD to the nearest 10^-4^ m.

### Statistical analyses

All analyses were performed using R 4.5.2 (R Core Team 2025). To examine the effects of ploughing (hypothesis a) and wildflower planting age (hypothesis c) on the number of overwintering bee nests, we fitted a generalized linear mixed model in the ‘glmmTMB’ package (Brooks et al. 2017) using a Poisson distribution, with the number of nests per plot summed across all sampling rounds within each sampling year as the response variable. Fixed effects included ‘ploughing’, ‘age’, their interaction to allow the ploughing effect to vary with wildflower planting age, and ‘sampling year’ (Suppl. S1a). ‘Age’ was defined as the number of years since the initial ploughing and sowing that occurred to establish the wildflower planting. ‘Plot’ was included as a random effect to account for random variation and repeated observations across sampling years. Sampling duration (number of days of continuous trap deployment) was not included as a covariate because sampling extended beyond the main emergence period of overwintering bees in all plantings.

Model fit was assessed using residual diagnostics of the ‘DHARMa’ package (Hartig 2024) as well as likelihood-ratio-tests of the ‘stats’ package (Lewis et al. 2011; R Core Team 2025). Model performance was assessed using marginal R^2^ from the ‘performance’ package (Lüdecke et al. 2021; Nakagawa et al. 2017). For the post-hoc analysis, estimated marginal means (EMMs) were obtained for different treatments (unploughed vs ploughed) and ages (0-4 years) using the ‘emmeans’ package (Lenth 2017; Searle et al. 1980), and pairwise contrasts between the EMMs were calculated to compare predicted nest counts between unploughed and ploughed plots across different wildflower planting ages. Percentage changes were derived from these contrasts on the response scale to quantify relative differences. For visualization purposes, model predictions were upscaled to estimate nest density (nest numbers per hectare).

To test whether smaller bees were more affected by ploughing than larger bees (hypothesis b), we used a permutation-based analysis of covariance in the ‘permuco’ package (Frossard & Renaud 2024), which does not rely on parametric ANOVA assumptions and was therefore appropriate because model residuals deviated from normality. Mean intertegular distance (ITD) was calculated for each plot and sampling year and used as the response variable. The model included ‘ploughing’ as a fixed factor and ‘age’ as a continuous covariate. The effect of ploughing was tested while controlling for differences in wildflower planting age using 2000 permutations.

## Results

We collected a total of 121 wild bees (2023: 56 bees; 2024: 65 bees) from wildflower plantings aged 0-4 years. Bees were collected in 74 out of the 368 e-traps, corresponding to a trapping success rate of 20.1 %. Bees included 27 species (see Table S2) belonging to the genera *Lasioglossum, Andrena, Bombus, Halictus* and *Melitta*. More than half of the bees could be classified as having emerged from overwintering (68 out of 121 wild bees = 56 %, see Table S2), and were linked to 63 nests. All other bees were excluded from further analysis because they were classified either as having emerged from an active nest or a nest built in the same season (17 bees = 14 %), or as ‘unknown’ (36 bees = 30 %).

Bee nest numbers were negatively affected by ploughing (estimate: −3.34, z = −2.49, *p* = 0.01, R^2^_*marginal*_ = 0.35; for coefficient table see table S1a), with the strongest reduction one year since establishment. Ploughing reduced nest numbers by −81.2 % at this age (EMM contrast for ploughed and unploughed plots one year since establishment: SE: 9.8 %, z = −3.22, *p* = 0.001). No effect of ploughing was detected two or three years since establishment (EMM contrasts for ploughed and unploughed plots: two years since establishment: *p* = 0.15; three years since establishment: *p* = 1.00). Predicted nest numbers in ploughed plots remained constant at the same low level (~ 150 nests per ha) across wildflower planting ages, but the interaction between age and ploughing was not significant (interaction term: estimate = 0.84, z = 1.58, *p* = 0.12).

Following wildflower planting establishment, nest numbers increased 5.2-fold in the first year since establishment without further ploughing (EMM contrast for unploughed plots one year since establishment vs. ploughed plots in the year of establishment: + 419.2 %, SE 401.0 %, z = 2.13, *p*= 0.03), resembling the development on perennial wildflower plantings (see Fig. 1). Thereafter, nest numbers in unploughed plots decreased with wildflower planting age (estimate = −0.86, z = −2.40, *p* = 0.02). Overall nest numbers were higher in 2024 than in 2023 (estimate = 0.89, z = 1.96, *p* = 0.05).

**Figure 1.**
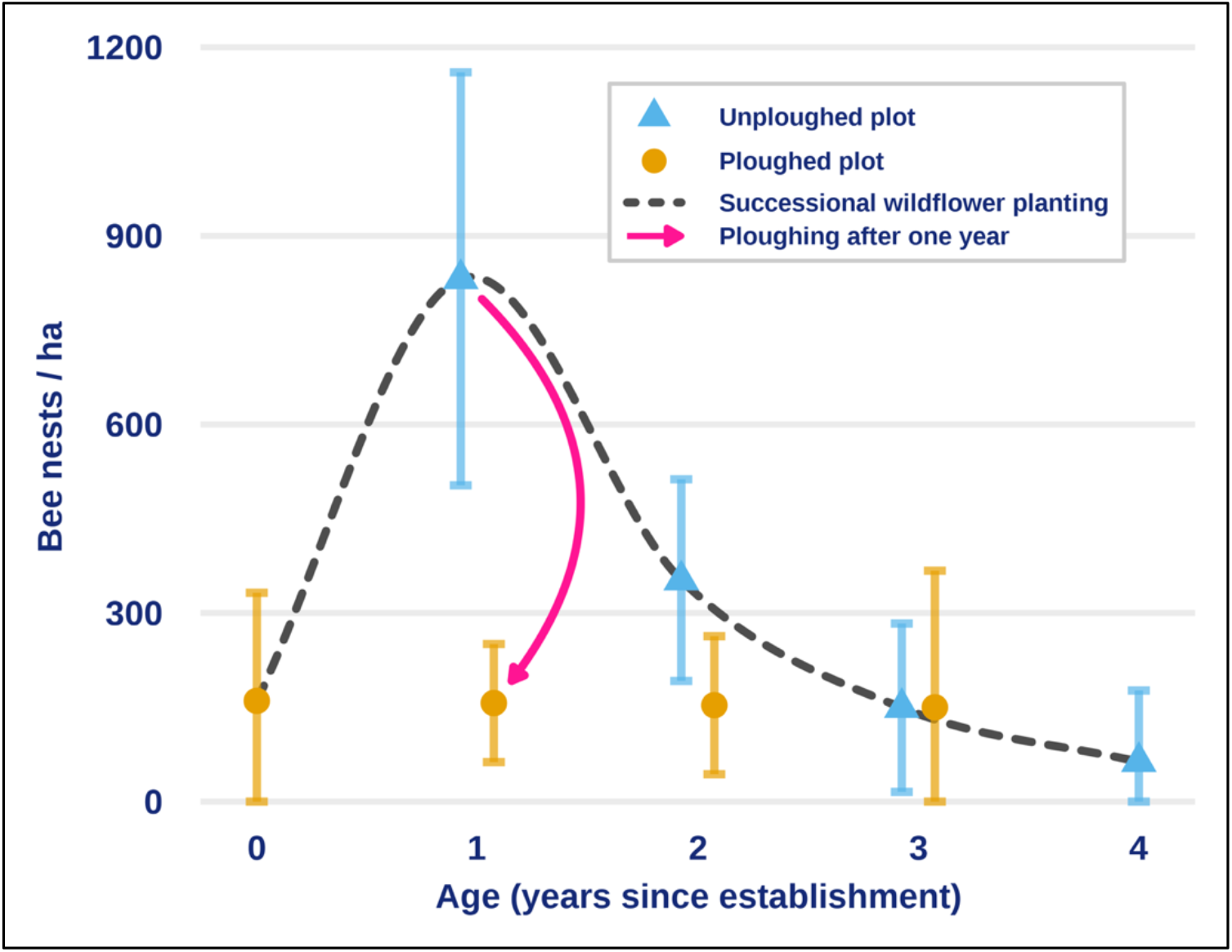
Estimated bee nest numbers per hectare on ploughed (triangles) and unploughed (points) plots 0-4 years since establishment; error bars indicate standard error of the means. Ploughing after one year (pink arrow) corresponds to the typical management in annual wildflower schemes and had the greatest negative effect on nest numbers. The dashed line represents an interpolation through model-predicted values to illustrate the temporal development of bee nest numbers in typical successional wildflower plantings (i.e., perennial wildflower schemes) which are initially ploughed for establishment and thereafter left unploughed. Nests were inferred from bees emerging from overwintering and were thus established in the preceding year.

The intertegular distance (ITD) of the 68 bees that were classified as having emerged from overwintering ranged from 0.7 mm (*Lasioglossum lucidulum*) to 5.6 mm (*Bombus terrestris*) (ITD mean: 1.43 mm, median: 1.00 mm). Bee body size did not differ between ploughed and unploughed plots (F_1,18_ = 0.25, permutation *p* = 0.67, Fig. S3a), nor did wildflower planting age affect bee body size (F_1,18_ = 0.02, permutation *p* = 0.88), although there was a visual tendency of ITD increasing with age (Fig. S3b).

## Discussion

We assessed nest numbers of overwintering ground-nesting wild bees in wildflower plantings. Plantings were either newly established or one to four years old, and within plantings unploughed control plots were compared with treatment plots ploughed in early spring. Ploughing reduced the number of nests from which bees successfully emerged, with the strongest effect one year since wildflower planting establishment. On unploughed control plots, bee nest numbers were highest after the first flowering season and declined thereafter, indicating highest habitat suitability for nesting in the year of establishment. In contrast, ploughed plots remained at consistently low nest numbers independent of how old they were prior to ploughing. We conclude that newly established wildflower plantings can rapidly become attractive nesting and overwintering habitat for ground-nesting bees, but that perennial plantings may become less attractive for nesting and overwintering as they get older. We further conclude that ploughing wildflower plantings after one flowering season, which corresponds to the typical management of annual wildflower plantings, creates an ecological trap for ground-nesting bees by likely increasing mortality following ploughing. Postponing annual ploughing of wildflower plantings, and potentially also of other crops, therefore has strong positive effects for wild bee conservation.

### Ploughing negatively affects bees nesting and overwintering in the ground

Our study is among the first experimental studies to provide causal evidence of the effects of ploughing on the entire bee community present in the soil. Although negative effects of ploughing have been previously suggested, these have mostly been based on associations between tillage intensity and bee occurrence or community composition (Shuler et al. 2005; Williams et al. 2010). The few existing experimental studies have mainly focused on single crop-associated species (Ullmann et al. 2016). Our findings therefore add to the still very limited evidence base showing that soil disturbance can reduce the successful emergence of ground-nesting bees at community level (Cuminale et al. 2026). This is particularly relevant because most wild bees nest below ground, whereas bee conservation mostly targets aboveground activity, for example enhanced foraging activity by increasing floral resources (Christmann 2022; Orr et al. 2022).

The observed reduction of nest numbers after ploughing may be explained by direct mortality or by physical damage to nests and brood cells (Tschanz et al. 2024). Mouldboard ploughing turns over the upper soil layer and therefore affects the nesting layer of many ground-nesting bee species (Harmon-Threatt 2020). Mortality may result from mechanical destruction of brood cells, exposure of overwintering stages to unsuitable abiotic conditions, or the collapse of nest tunnels above brood cells (Shuler et al. 2005). Nonetheless, some bees still emerged from ploughed plots, indicating that ploughing is not necessarily lethal to all individuals. By fitting a supplementary presence-absence model, we could confirm that only the nest numbers, but not the probability of finding a nest was influenced by ploughing (Suppl. S1b). Survival may depend on nest depth, brood-cell structure, soil texture, moisture conditions, or the precise position of brood cells (Tschanz et al. 2024). While a generally disproportionate effect of ploughing on larger insects has been reported (Hellerich et al. 2026), we found no evidence that bee size influenced survival. Nesting depth appears to be unrelated to body size but may vary among species (Cane 1991), explaining why species of varying body size survived. This is in line with cases where several individuals of the same species emerged from the same nest on our plots. Alternatively, some brood cells could have withstood the mechanical disturbance caused by ploughing. The absence of a body-size effect should nevertheless be interpreted cautiously, because only a few individuals emerged from ploughed plots.

### Age of wildflower plantings has non-linear effects on bee nest numbers

We expected unploughed wildflower plantings to accumulate bee nests over time since establishment because perennial wildflower plantings provide continuous, undisturbed habitat over multiple years (Albrecht et al. 2020; von Königslöw et al. 2022) and bees are known for philopatry (Harmon-Threatt 2020). Instead, we observed the highest nest numbers one year after planting establishment, meaning that wildflower plantings were most strongly used for nesting and overwintering during their first flowering season. Although the supplementary presence-absence model showed that in general wildflower plantings are used as nest sites independent of age (Suppl. S1b), our results suggest that perennial wildflower plantings are less attractive as nesting habitat for ground-nesting wild bees with increasing age as nest numbers did not increase but decreased with planting age.

Nesting habitat suitability may generally depend on a combination of floral resources and ground conditions (Sarthou et al. 2014), both of which favour nesting: flowers provide nearby pollen and nectar resources for central-place foraging bees, while bare ground facilitates nest construction (Antoine & Forrest 2021; Cane 1991; Tsiolis et al. 2022; Zurbuchen et al. 2010). Our findings suggest particularly high nesting suitability in the year of wildflower planting establishment. In this year, sites initially underwent ploughing for seedbed preparation, which created open ground. Supplementary flower-cover assessments indicated that the plants sown to establish the planting only started flowering in June (Suppl. S4; Fig. S4). Even as flower cover increased in later months in the year of establishment, soil conditions remained relatively open because dead plant material and litter were still largely absent (pers. obs.). Most bees collected in emergence traps after the first winter emerged in April and May (Fig. S4). These bees had nested during the preceding year of establishment. As nest building soon follows emergence, and flowering in the year of establishment only started in June, habitat suitability for some species may therefore have been determined more by ground conditions than by the presence of flower resources.

Decreasing nest numbers with increasing wildflower planting age may be linked to decreased nesting habitat suitability, even though sites remain undisturbed. This may be explained by vegetation succession, leading to denser vegetation cover, reduced bare ground, and increased dominance of grasses (Frank et al. 2012; Ganser et al. 2019; Pywell et al. 2011). Our findings align with the limited existing evidence on these nesting dynamics. Boetzl et al. (2022) found the highest richness and activity density (a measure for abundance) of ground-nesting bees in flowering fields one year after the last soil disturbance compared with both recently disturbed and older sites. Williams et al. (2024) also reported decreasing nest numbers with increasing wildflower planting age, although their sampling approach may have captured active rather than overwintering nests, and different factors may influence the selection of sites for reproduction and overwintering (Yanega 1990). Resulting lower nest numbers may, in turn, reduce the proportion of ground-nesting bees in the foraging bee community (Steffan-Dewenter & Tscharntke 2001). By contrast, the effectiveness of wildflower plantings for crop pollination can increase with age, likely because local pollinator populations require time to build up (Albrecht et al. 2020; Blaauw & Isaacs 2014) and because floral diversity may increase in the second year of establishment (von Königslöw et al. 2022). However, without interventions such as adapted mowing, pollinator diversity is likely to saturate or decline, driven by decreasing floral diversity in older plantings (Blaauw & Isaacs 2014; von Königslöw et al. 2022). Together with our findings, this suggests that wildflower planting age may affect ground nesting and overwintering of bees differently than aboveground flower visitation, highlighting a trade-off between nesting and foraging resources: wildflower plantings may up to a certain age support pollinators through diverse floral resources and landscape connectivity, while their suitability as nesting habitat for ground-nesting bees declines with age from the outset. Our results should therefore not be interpreted as an argument against perennial wildflower plantings, but as a call for a mosaic of different successional stages (Eccard 2022; Ganser et al. 2019).

### Annual wildflower plantings are ecological traps for ground-nesting wild bees

Our results support the conclusion that annually ploughed wildflower plantings act as ecological traps for ground-nesting wild bees. Individuals may select newly established wildflower plantings as habitats of high apparent nesting quality, but annual wildflower plantings are commonly ploughed again after one flowering season and before bee emergence in the following spring. Their management cycle thus overlaps with the bee life cycle. Ploughing both creates conditions suitable for nesting and at the same time increases mortality after individuals have selected sites for nesting, resulting in wildflower plantings acting as ecological traps (Robertson & Hutto 2006). Ecological trap-effects likely differ among insect taxa because of variation in phenology, overwintering location, dispersal ability, sensitivity to soil disturbance or development dynamics on wildflower plantings (Frank et al. 2012; Ganser et al. 2019; Haaland et al. 2011; Hellerich et al. 2026). For example, it was not observed for epigean arthropods in annually ploughed wildflower plantings (Füglistaller et al. 2018).

The relative attractiveness of wildflower plantings as nesting habitat still remains unknown, and our experiment does not test habitat preference relative to all available alternative habitats in the surrounding landscape such as arable fields, field margins, bare tracks or other semi-natural habitats. However, our data show that bees prefer wildflower plantings in the year of their establishment for nesting and overwintering, and that ploughing strongly reduces nest numbers, and thus fitness. Even without a choice experiment, we can thus conclude that annually ploughed wildflower plantings can act as ecological traps for the bees that use them for nesting, because identifying an ecological trap concerns the consequences of habitat selection behaviour rather than population-level effects (Robertson & Hutto 2006). This risk is likely to be underestimated when habitats are evaluated only by their foraging quality (e.g., flower cover or pollinator visitation during the flowering season), because the life stage of bees in the ground is immobile and present long after the aboveground foraging period, which may only last a few days. Ploughed crop fields adjacent to flower resources may also be similarly attractive as nesting habitat as wildflower plantings in their first year because of open ground conditions. To what extend crop fields are active sinks for ground-nesting wild bees requires further testing which should include factors influencing overwintering in ground-nesting bee species, as overwintering habitat preferences and species-specific behaviours are still mostly unknown (Westrich 2018)

Our findings underline the importance of treating annual and perennial wildflower plantings as functionally distinct habitats for wild bee conservation. Annual wildflower plantings can provide temporary floral resources and may have biodiversity value in their own right (Westbrook et al. 2024), but they are not a substitute for perennial habitat when the goal is to support the full life cycle of ground-nesting wild bees. They may even undermine their own conservation objective by attracting bees to nesting habitat that is ploughed before successful emergence, thereby drastically reducing reproductive success and creating an ecological trap. Meanwhile, our results also indicate that perennial wildflower plantings lose attractiveness as nesting habitat for ground-nesting bees at later successional stages, highlighting potential for management optimization. Current agri-environmental schemes for wildflower plantings mainly emphasize flower availability, visual attractiveness and pollinator activity (Schmidt et al. 2022), but to support ground-nesting bees effectively, they should also account for their requirements to nesting and overwintering resources (Christmann 2022), for example by maintaining different successional stages to create partial open ground (Eccard 2022; Tsiolis et al. 2022) or by reducing management intensity through shallow ploughing or conservation tillage (Stinner & House 1990). The high nest numbers in early-successional wildflower plantings indicate that soil conditions may be a major determinant of nesting habitat suitability and highlight a management trade-off: ploughing can create attractive open ground where vegetation succession and litter may have reduced habitat suitability, but it can also destroy nests in the ground and prevent successful overwintering and reproduction. Especially annual soil management may therefore have unexpectedly large ecological consequences for ground-nesting wild bees in agricultural landscapes.

## Acknowledgements

We gratefully acknowledge funding from the Ministry for the Environment, Climate and Energy Sector of Baden-Württemberg. We thank the lower nature conservation authorities of Freiburg and Emmendingen for issuing the required permits, as well as the Regierungspräsidium Freiburg (Ref. 56 and 33), NABU Freiburg and all involved farmers for their support in selecting and managing the study sites.

## Notes

### Competing Interest Statement

The authors have declared no competing interest.

