## Supplementary information for "Annual ploughing turns wildflower plantings into ecological traps for ground-nesting wild bees"

*shared last authorship

^1^ Nature Conservation and Landscape Ecology, University of Freiburg, Stefan-Meier-Straße 76, 79104 Freiburg, Germany

^2^ Centre for Agri-Environmental Research, University of Reading, Earley Gate, Reading RG6 6AR, United Kingdom

^3^ Animal Ecology, Department of Biology, University of Marburg, Karl-von-Frisch-Straße 8, 35043 Marburg, Germany

This document contains:

**S1: Model terms and coefficients**	**2**

S1a: Nest-count model	2

Model term	2

Model coefficient table	2

S1b: Supplementary binomial ‘presence-absence’ model	3

Model term	3

Results	3

Model coefficient table	3

**S2: Emergence-trapped bees**	**4**

**S3: Bee body size across treatments and age**	**5**

**S4: Supplementary flower cover assessment**	**6**

Method	6

Results	6

References	7

### S1: Model terms and coefficients

#### S1a: Nest-count model

##### Model term

nest_count_model <- glmmTMB(

num_nests ~ age * ploughed + sampling_year + (1 | plot),

family = poisson)

##### Model coefficient table

**Table S1a: Nest model coefficients**

|  |  |  | **num_nests** |  |  |
| --- | --- | --- | --- | --- | --- |
| Predictors | | *Log-Mean* | *std. Error* | *Statistic* | *p* |
| (Intercept) | | 1.65 | 0.92 | 1.79 | 0.073 |
| age | | -0.86 | 0.36 | -2.40 | **0.017** |
| ploughed [1] | | -3.34 | 1.34 | -2.49 | **0.013** |
| sampling year [2024] | | 0.89 | 0.45 | 1.96 | **0.050** |
| age × ploughed [1] | | 0.84 | 0.53 | 1.58 | 0.115 |
| **Random Effects** | | | | | |
| σ^2^ | | 0.50 | | | |
| τ_00_ _plot_ | | 1.12 | | | |
| ICC | | 0.69 | | | |
| N _plot_ | | 28 | | | |
| Observations | | 46 | | | |
| Marginal R^2^ / Conditional R^2^ | | 0.353 / 0.799 | | | |

#### S1b: Supplementary binomial ‘presence-absence’ model

We fitted an additional ‘presence-absence’ model to test whether ploughing and wildflower planting age influenced the probability of bee nest occurrence rather than nest number, using a binomial distribution. This model included the same fixed and random effects as the nest-count model, with nest occurrence (yes / no) as the response variable. This allowed us to distinguish effects on the probability of detecting at least one nest from effects on overall nest numbers.

##### Model term

nest_presence_model <- glmmTMB(

nest_obs ~ age * ploughed + sampling_year + (1 |plot),

family = binomial)

##### Results

This binomial presence-absence model indicated that neither ploughing (estimate = -3.99, z = -1.29, *p* = 0.20, R^2^*_marginal_* = 0.23) nor wildflower planting age (estimate = -0.87, z = -1.11, *p* = 0.27) had a significant effect on the probability of bee nest occurrence.

##### Model coefficient table

**Table S1b: Binomial nest model coefficients**

|  |  |  | **nest_obs** |  |  |
| --- | --- | --- | --- | --- | --- |
| Predictors | | *Log-Odds* | *std. Error* | *Statistic* | *p* |
| (Intercept) | | 2.43 | 2.29 | 1.06 | 0.290 |
| age | | -0.87 | 0.78 | -1.11 | 0.265 |
| ploughed [1] | | -3.99 | 3.08 | -1.29 | 0.195 |
| sampling year [2024] | | 0.90 | 0.89 | 1.01 | 0.312 |
| age × ploughed [1] | | 0.75 | 0.98 | 0.76 | 0.446 |
| **Random Effects** | | | | | |
| σ^2^ | | 3.29 | | | |
| τ_00_ _plot_ | | 1.68 | | | |
| ICC | | 0.34 | | | |
| N _plot_ | | 28 | | | |
| Observations | | 46 | | | |
| Marginal R^2^ / Conditional R^2^ | | 0.234 / 0.493 | | | |

### S2: Emergence-trapped bees

**Table S2: Total number of bees collected in the emergence traps, number of bees classified as overwintered, and number of overwintering nests per species and genus.**

| **Species** | **Total number** | **Number of bees classified as overwintered** | **Number of overwintering nests** |
| --- | --- | --- | --- |
| ***Andrena*** | **33** | **9** |  |
| *alfkenella* | 1 |  |  |
| *bicolor* | 1 |  |  |
| *dorsata* | 9 | 3 | 1 |
| *flavipes* | 2 |  |  |
| *minutula* | 7 | 4 | 2 |
| *minutuloides* | 9 | 2 | 1 |
| *ovatula* | 1 |  |  |
| *rosae* | 2 |  |  |
| *scotica* | 1 |  |  |
| ***Bombus*** | **5** | **2** |  |
| *pratorum* | 1 |  |  |
| *terrestris* | 4 | 2 | 2 |
| ***Halictus*** | **3** | **1** |  |
| *eurygnathus* | 2 | 1 | 1 |
| *simplex* | 1 |  |  |
| ***Lasioglossum*** | **79** | **56** |  |
| *fulvicorne* | 4 |  |  |
| *glabriusculum* | 30 | 27 | 27 |
| *laticeps* | 2 |  |  |
| *leucozonium* | 8 | 3 | 3 |
| *lucidulum* | 1 | 1 | 1 |
| *malachurum* | 3 |  |  |
| *morio* | 2 | 2 | 2 |
| *pauxillum* | 2 |  |  |
| *punctatissimum* | 1 | 1 | 1 |
| *puncticolle* | 2 |  |  |
| *sabulosum* | 4 | 4 | 4 |
| *sexnotatum* | 2 | 2 | 2 |
| *zonulum* | 18 | 16 | 16 |
| ***Melitta*** | **1** |  |  |
| *leporina* | 1 |  |  |

### S3: Bee body size across treatments and age

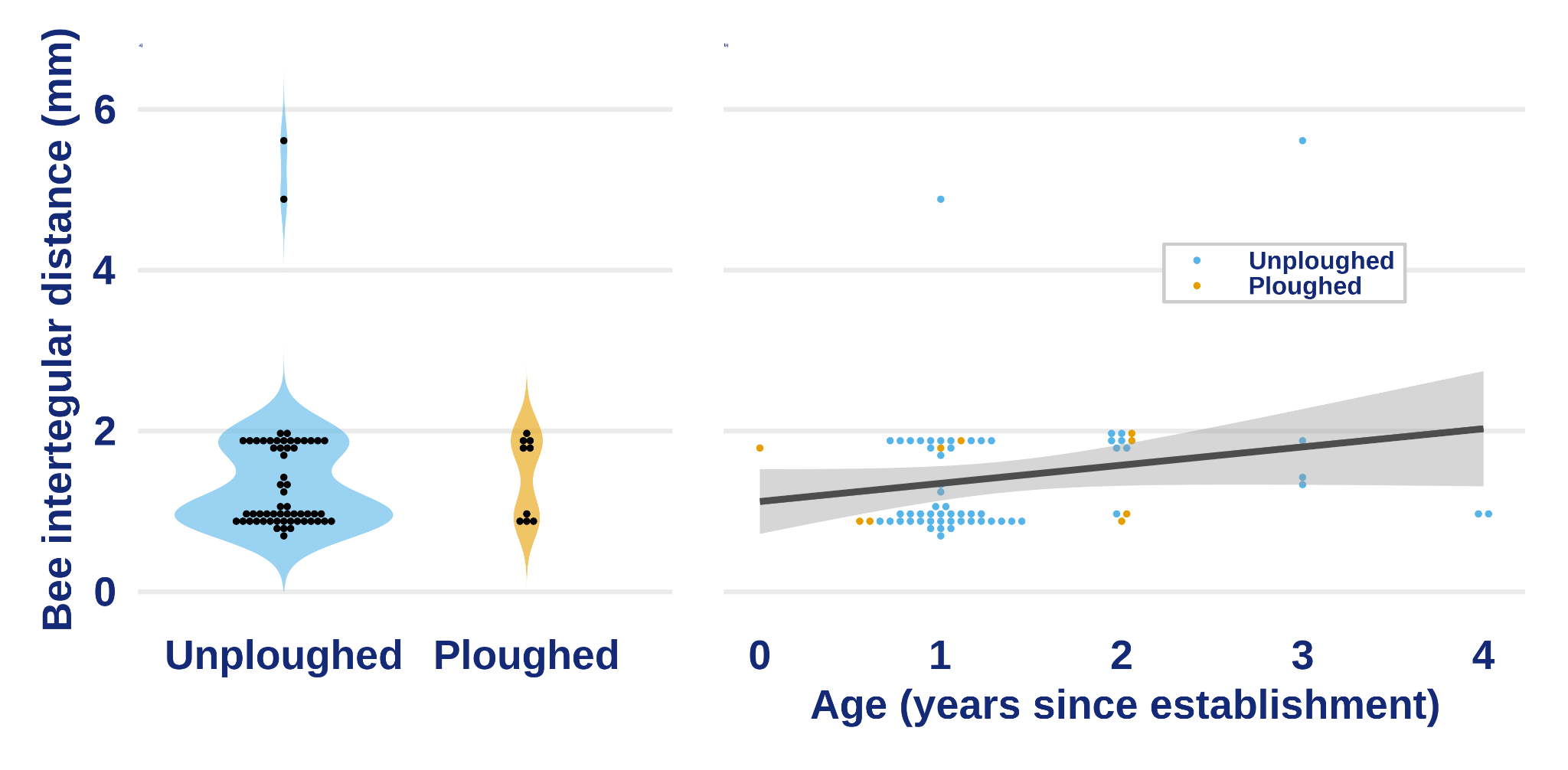

**Figure S3: Individual bee intertegular distances (points) across a) treatments (unploughed vs. ploughed), and b) years since wildflower planting establishment. The grey line represents a linear model fitted to the data for visualisation; ribbons indicate 95 % confidence intervals.**

### S4: Supplementary flower cover assessment

#### Method

We quantified flower cover on newly established as well as 1-4 year old wildflower plantings. Surveys took place once per month from April to August in 2023 and 2024. The only exception was in 2023, when surveys on 1-3 year old plantings began in May.

During each survey round, two 1 m^2^ squares were selected per planting to represent the flowering plant community present at the site as completely as possible (Blaydes et al., 2024). Within each square, the percentage cover of each flowering plant species was estimated separately in 5% increments. Species represented by only a single flower were assigned a cover value of 1%. Flower cover per square was calculated as the sum of the cover values of all flowering plant species and could therefore exceed 100% where flowers of different species overlapped vertically. Total mean flower cover per planting was then calculated by taking the mean of the two quadrats.

To visually assess the seasonal development of flower cover in relation to bees emerging from nests, we plotted the raw flower-cover data for wildflower planting age 0-4 together with the number of overwintered wild bee nests inferred from bees sampled with emergence traps in the following year. Seasonal developments were illustrated using third-order polynomial (cubic) regressions.

#### Results

In total, 104 flower-cover surveys were conducted. Total flower cover ranged from 0% to 132%, with a mean of 57.9% (standard deviation = 29.5%). During the year of establishment (Age = 0 years), flower cover averaged 30.2% (standard deviation = 34.9%; range: 0-100%; n = 20). In 1 year old plantings, mean flower cover increased to 68.5% (standard deviation = 19.4%; range: 15-104%; n = 33); in 2 year old plantings it averaged 58.6% (standard deviation = 19.4%; range: 2.5-92.5%; n = 23), which barely changed in three year old plantings, where flower cover averaged 58.4% (standard deviation = 23.7%; range: 15-100%; n = 18). In 4 year old plantings mean flower cover increased to 75.6% (standard deviation = 36.3%; range: 12.5-132%; n = 10). Newly established wildflower plantings did not show measurable flower cover before June, whereas older plantings always showed flower cover in April already (Fig. S4). Regardless of time since establishment, flower cover always increased as the season progressed.

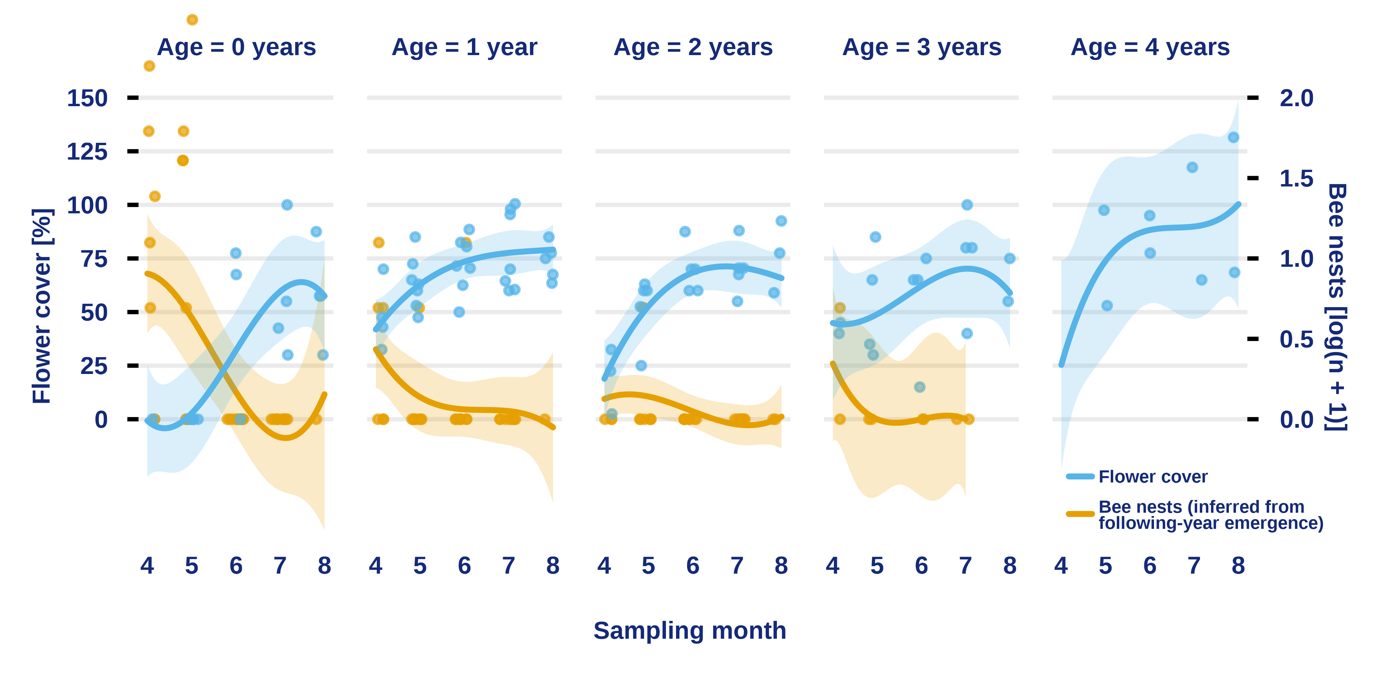

**Figure S4: Seasonal development of flower cover in wildflower plantings aged 0-4 years (blue) and the number of ground-nesting bee nests established during the corresponding nesting season (orange). Flower cover was recorded during each nesting season, whereas nests were inferred from overwintered bees captured with emergence traps in the following year. Thus, flower-cover and nest data within each panel refer to the same nesting season. No bees were sampled after the fifth flowering season (Age = 4 years) and no nesting can thus be inferred for this season. Blue points show raw total flower cover per planting, and orange points show the raw number of inferred overwintered bee nests, expressed as log(n+1). Lines show fitted third-order polynomial (cubic) regressions across sampling months and ribbons indicate 95% confidence intervals.**
